# Systematic shifts in the variation among host individuals must be considered in climate-disease theory

**DOI:** 10.1101/2021.08.30.458260

**Authors:** Joseph R Mihaljevic, David J. Páez

## Abstract

To make more informed predictions of host-pathogen interactions under climate change, studies have incorporated the thermal performance of host, vector, and pathogen traits into disease models. However, this body of work has ignored the fact that disease spread and long-term patterns of host population dynamics are largely determined by the variation in susceptibility among individuals in the host population. Furthermore, and especially for ectothermic host species, variation in susceptibility is likely to be plastic, influenced by variables such as environmental temperature. For example, as host individuals respond idiosyncratically to temperature, this could affect the population-level variation in susceptibility, such that there may be predictable functional relationships between variation in susceptibility and temperature. Quantifying the relationship between temperature and among-host trait variation will therefore be critical for predicting how climate change and disease will interact to influence host-pathogen population dynamics. Here, we use a model to demonstrate how short-term effects of temperature on the variation of host susceptibility can drive epidemic characteristics, fluctuations in host population sizes, and probabilities of host extinction. Our results emphasize that more research is needed in disease ecology and climate biology to understand the mechanisms that shape trait variation, not just trait averages.

## Introduction

Theory suggests that climate warming can destabilize natural population fluctuations, e.g., moving from stable equilibria or cycles to unstable cycles or chaos [2, 3, 4, 5]. Such destabilization can in turn drive population declines or extinctions, with auxiliary effects that ripple through an ecosystem [6, 7]. Independent of climate, pathogens that cause mortality can also drive periodic fluctuations in population size (e.g., cycling), that can in turn lead to host population decline, local extirpation, or even species extinction [8, 9, 10]. Understanding how climate and host-pathogen interactions combine to determine population declines over the long term is however exceptionally challenging [11, 12, 13]. While there is concern that climate change will exacerbate disease processes such as the frequency, duration and magnitude of epidemics, the mechanisms driving such effects are still poorly characterized, such that our ability to predict climate’s future effects on host-pathogen interactions is unclear [14]. To improve our mechanistic understanding, there has been a surge in modeling studies that incorporate the effects of temperature on the plastic traits of hosts and pathogens that drive transmission dynamics. Specifically non-linear thermal performance curves (TPCs, also known as thermal reaction norms) have been incorporated into mechanistic, mathematical models of host-pathogen (or host-vector-pathogen) interactions [15, 16, 17], which has clarified the specific mechanisms by which temperature drives spatial and temporal patterns of pathogen prevalence in a variety of systems [18, 19]. For example, accounting for the temperature-dependence of traits that influence mosquito demography and pathogen transmission has led to well-validated predictions of geographic locations expected to have the highest prevalence of certain mosquito-borne diseases [15, 20, 21, 22, 23]. While much progress has been made to understand the mechanistic effects of temperature using TPCs and modeling, we argue that the research has largely ignored variation in the shapes of reaction norms among host individuals. As we describe, accounting for how temperature systematically changes the variability in traits expressed among host individuals is likely essential for understanding the ecological outcomes of host-pathogen interactions under climate change.

The variation in trait values among individuals (e.g., transmission rates) has fundamental consequences on disease dynamics [24, 25], yet the vast majority of research on thermal performance curves in disease ecology has focused on population-level trait averages. Yet, the variability in plastic trait values among host individuals may also systematically shift along a thermal gradient (Fig. 1), but there is no current theory to understand the implications of temporally shifting trait variation on disease dynamics. In the current literature on temperature-dependent disease ecology, “trait variation” has been used very specifically: the models that integrate TPCs assume that the average values of traits within a population vary as functions of temperature (e.g., solid lines in Fig. 1), and that all individuals share the same trait value at any value of temperature. In other words, “trait variation” is taken to mean that trait averages vary across a temperature gradient. However, because hosts respond individualistically to temperature, we must not only be concerned about population-level trait averages, but also with the variation in trait values among individuals. “Trait variation” can therefore also reference the fact that trait values vary among host individuals, as emphasized by Cator et al. [18]. However, there has been almost no research on how among-host variability in trait values shifts across temperature gradients. In fact, the among-host variability in trait values may have predictable, functional relationships with temperature that are independent from the relationships between temperature and the average, population-level trait values (Fig. 1). Relationships between temperature and among-host trait variation will very likely have meaningful impacts on short- and long-term disease dynamics that should not be ignored.

**Figure 1:**
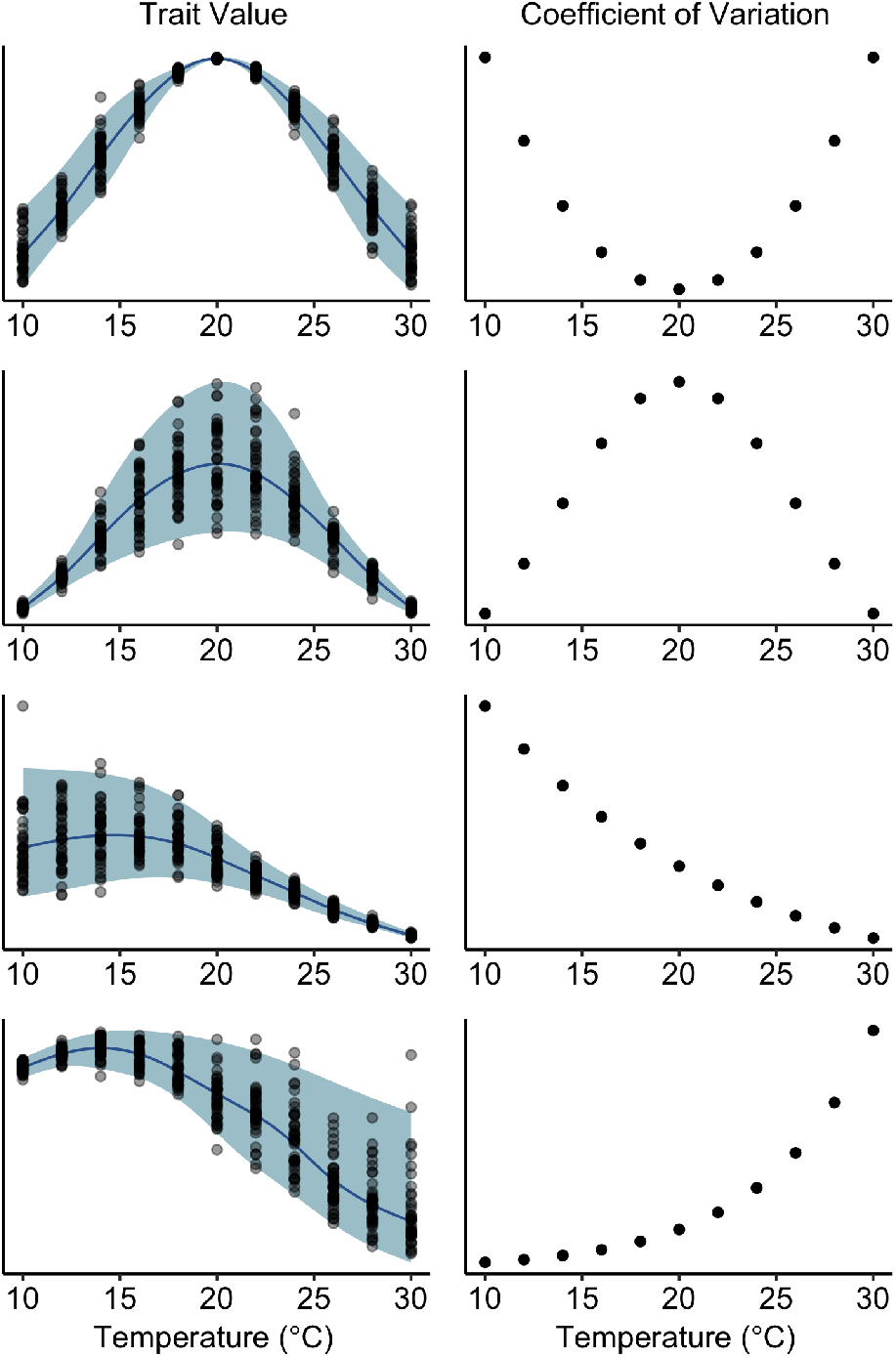
Simulated thermal performance curves in which variability among individuals systematically shifts across the temperature gradient. The left column shows trait values measured across 40 individuals per temperature (points), and the right column shows the coefficient of variation of the trait value among individuals at each temperature. The line is the median (with 95% confidence ribbon) of an additive quantile regression model fit using the R package qgam [1].

In this study, we provide a structured argument for why we need more empirical and theoretical research related to the effects of temperature on among-host variation in host and vector traits relevant for disease transmission. First, we provide a brief overview of selected literature that collectively demonstrate that temperature does affect among-host trait variation in several ectotherm vector and host systems, supporting that such patterns are likely widespread. Second, we provide theoretical background on why among-host trait variation is important for disease epidemics and long-term stability of host populations that are infected by pathogens that affect survivorship. Specifically, we use a host-pathogen model to demonstrate that if temperature affects the variation in susceptibility among host individuals, our theoretical expectations for host population stability can change drastically, for example, moving from stable to unstable cycles. We believe that this analysis, while simple, reveals that our understanding of how climate and disease will combine to affect long-term host-pathogen dynamics may be flawed without considering effects of how temperature alters patterns in among-host trait variation. This has implications for projecting the long term impacts of infectious disease on vulnerable host populations, especially in a changing climate.

## Methods

### A brief review

Across the literature on thermal ecology (even beyond thermal disease ecology), there has been very little focus on empirically measuring whether temperature affects among-individual variation. To visually illustrate our definition of trait variation, Figure 1 illustrates hypothetical examples of how trait variation can change across temperature gradients, independent of trait averages. While we did not attempt a systematic review of the vast thermal ecology literature (e.g., meta-analysis), we did find a few anecdotal data sets that support several of the hypothetical relationships in Figure 1, and others. For example, Careau et al. [26] found that the mean jump time and distance jumped until exhaustion in clawed frogs (*Xenopus tropicalis*) declined across a temperature gradient, and so did the among-host variation in these traits (see Careau et al. [26]’s Fig. 1c,d, similar to our Fig. 1, third row). Young and Gifford [27] measured the swimming speed of salamanders (*Desmognathus brimleyorum*) across a temperature gradient, finding a unimodal relationship for the mean, and a higher variation at the peak performance temperature compared to the thermal limits (see Young and Gifford [27]’s Fig. 2, similar to our Fig. 1, second row). The mean maximal sprint speed of common lizards (*Zootoca vivipara*) increased with temperature, and so did the variation among individuals (see Artacho et al. [28]’s Fig. A1, similar to our Fig. 1, last row, but with an increasing mean).

**Figure 2:**
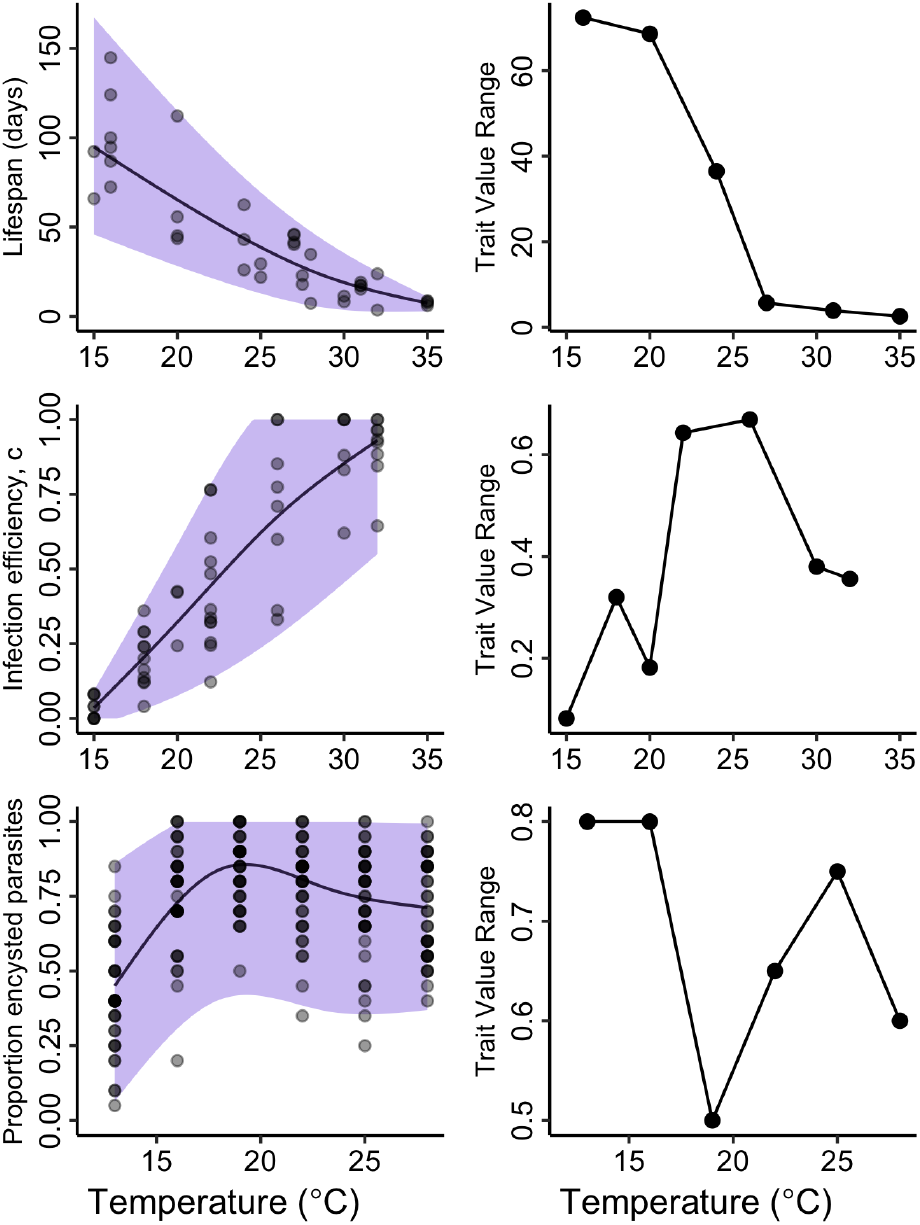
Data on thermal performance from the literature in which variability among individuals systematically shifts across the temperature gradient. The left column shows trait values measured across individuals per temperature (points), and the right column shows the range (maximum - minimum) of the trait value among individuals at each temperature (only including temperatures with at least 3 data points). The line is the median (with 95% confidence ribbon) of an additive quantile regression model fit using the R package qgam [1]. The top two panels are data on *Cx. pipiens* [23], the middle panel is in reference to West Nile virus, and the lower panel is data on parasites encysting in frog tadpoles [29].

There are similar anecdotal patterns for vector traits that influence pathogen transmission. Shocket et al. [23] provide several TPCs of mosquito traits, and for some traits, variation among mosquito individuals seems to systematically shift across a thermal gradient For example, the mean lifespan of *Culex pipiens* declines with temperature and so does variation among host individuals (Fig. 2, top row); the mean infection efficiency of Cx. pipiens for West Nile virus increases with temperature, and the variation seems to be unimodal or generally increases with temperature (Fig. 2, middle row). The original data was not collected in a way to explicitly measure such variation, however, so the patterns are not well resolved for many traits.

In comparison with disease vectors, there are few data sets on the thermal performance of transmission-related traits in ectothermic hosts, making it difficult to generalize how trait variation scales with temperature. Some studies have estimated TPCs for host traits important for immune system dynamics [29, 30, 31, 32]. Yet, while these traits are likely linked to transmission, their specific relationship is unknown. Nevertheless, the data in [29] provide an example of how the variation in traits related to host-pathogen interactions changes with temperature. Overall, temperature had a quadratic effect on the likelihood of parasites encysting in the host, and the variation across individuals changed non-linearly as a function of temperature (Fig. 2, bottom row).

Interestingly Elderd and Reilly [33] explicitly measured the effect of temperature on transmission heterogeneity and used a model to project the effects on host population dynamics. The authors found that warmer temperatures decreased an insect host’s variation in susceptibility to a virus compared to colder temperatures, despite the fact that the average susceptibility was indistinguishable between a cold and a warm temperature. The decrease in variation was explained by an increase in the feeding rate of insects under warmer temperatures, which increases the likelihood of consuming virus particles present on foliage. Using a model similar to the one described below, the authors suggest that the reduced variation in susceptibility with increasing temperatures could result in larger epidemic sizes [33]. This study focused on only two temperature treatments, however, making it difficult to characterize any non-linear relationship between temperature and the variation in susceptibility among host individuals.

Overall, temperature likely has systematic effects on the variation of many traits across diverse host-pathogen systems, and such functional relationships are possibly system-specific. General patterns are however difficult to determine, because studies have overwhelmingly emphasized the effect of temperature on trait averages without explicitly considering variation in the effect of temperature across host individuals.

Across ecological systems, there is a rising appreciation that the degree to which traits vary among individuals can determine long-term patterns and large-scale processes in ecological time [34, 35, 36, 37, 38]. Indeed, a fundamental result from host-pathogen theory is that, when host individuals vary in traits such as susceptibility or pathogen shedding rate, the magnitude of variation (i.e., high or low variation) determines the size of disease outbreaks and the long-term cycling of disease in host populations [39, 40, 24, 41, 42] (Fig. 4). For example, models that allow for variation in host susceptibility demonstrate that hosts with high resistance and those with high susceptibility have strong effects on epidemic patterns and ultimately how many hosts become infected during an epidemic [25, 43, 44, 45, 46]. At the beginning of outbreaks, the presence of high-susceptibility hosts can lead to faster initial spread of the pathogen, but as the epidemic proceeds, high-resistance hosts generally slow the spread [45]. In fact, high-resistance hosts have disproportionate effects, such that small numbers of these resistant hosts cause epidemics to be smaller, especially in host populations of high density [43, 47, 48]. Moreover, if pathogen infection results in host mortality, even in a moderate fraction of cases, then variation in susceptibility determines how many hosts die during an outbreak (Figure 4). Variation in susceptibility therefore has long-term impacts on the cycling of host population abundances and the ultimate threat of pathogens to host population stability and extirpation [43, 49, 45].

**Figure 3:**
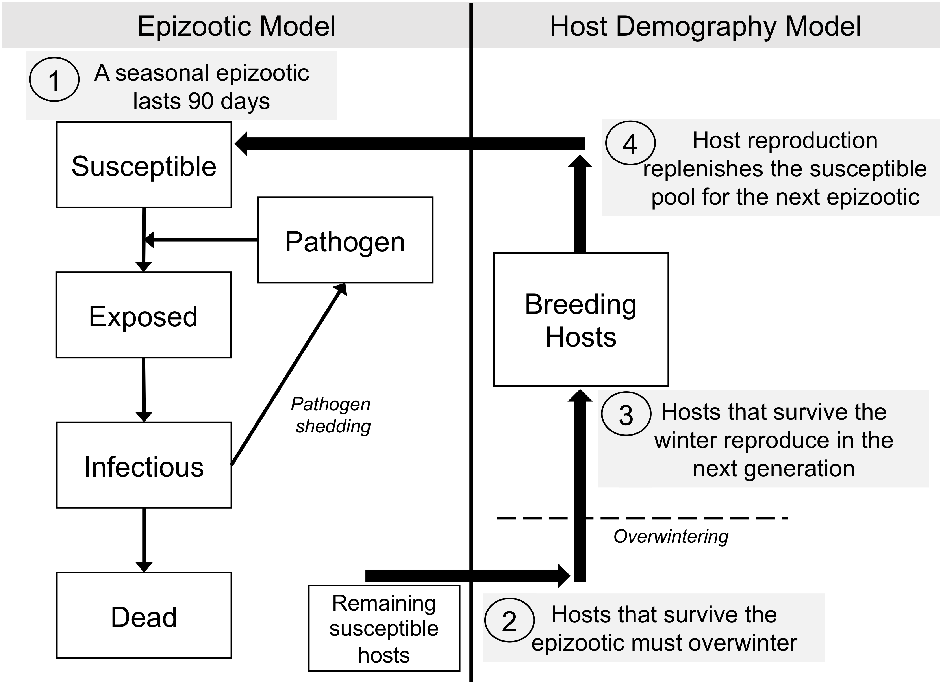
Schematic outlining the linkages between the epizootic and host demography models.

**Figure 4:**
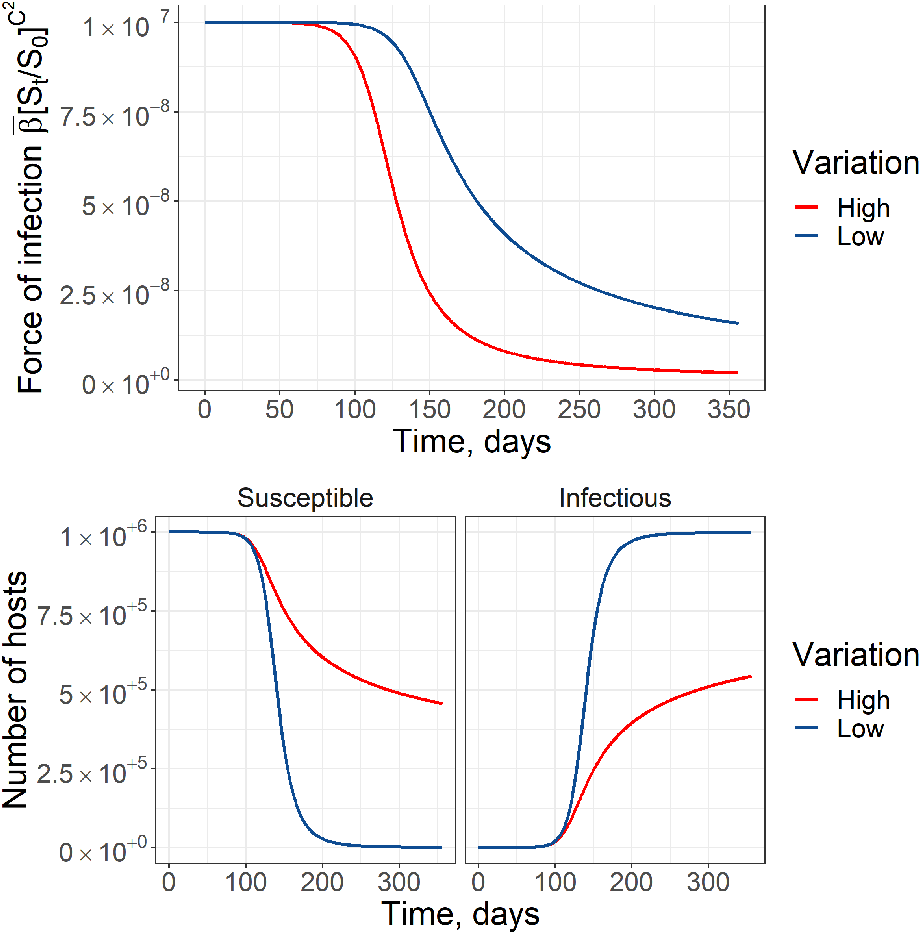
Epizootic dynamics are affected by heterogeneity in susceptibility. The top panel shows how the force of infection changes during an epizootic. The faster decline in the force of infection with high heterogeneity results in fewer hosts contracting infection, as shown in the bottom panels. This simulation was conducted using a compartmentalized Susceptible-Exposed-Infectious (SEI) model as described in the main text. 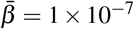, blue line *C*^2^ = 0.25, red line *C*^2^ = 5, *δ* = 1 *×* 10^*−*4^, Initial number of susceptible hosts, *S*_0_ = 1 *×* 10^6^.

This theory that describes how trait variation affects disease dynamics, however, assumes that the values of an individual host’s traits are fixed during its lifetime. When host traits related to transmission are plastic and depend on temperature, the assumptions of these previous models are violated and their results are no longer valid. This is problematic because for many host-pathogen systems, epidemics occur while climate is seasonally fluctuating [50, 51]. New theory is therefore needed to evaluate the implications of when variation of traits among host individuals shifts as temperature fluctuates during an epidemic.

### A mathematical model

Here we introduce a model to demonstrate how systematic variation in thermal performance curves among host individuals can drive epizootic dynamics and host population cycles. The model has two components. First, a compartmental disease model (i.e., epizootic model) describes how a pathogen transmits among a host species. Importantly, host individuals are assumed to vary in susceptibility to infection with the pathogen (i.e., some hosts are more resistant and some are more susceptible) [25, 46]. Second, we link this epizootic model with a simple model of host reproduction (i.e., host demography model) to understand how pathogen-induced mortality in the host population can influence long-term patterns in host population density [43, 52] (Fig. 3).

The epizootic model is described by the following set of differential equations, with state variables representing the densities of the host or pathogen:

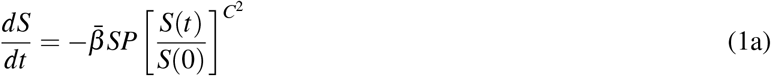

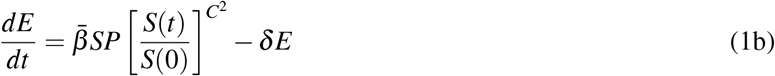

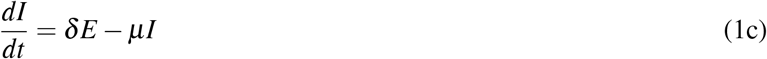

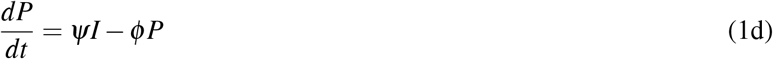

In this model, infectious hosts *I* shed the pathogen (*P*) into the environment at rate *Ψ*. Susceptible hosts (*S*) become infected through direct contact with the pathogen. Following a period of incubation, defined by *δ*, hosts exposed (*E*) to the pathogen become infectious. We assume that infection ultimately causes mortality at rate *µ*, and pathogens in the environment decay over time at rate *ϕ*.

The force of infection in this system ultimately determines how many hosts will die of the pathogen during an outbreak. Importantly here, we assume that the force of infection is affected by variability in susceptibility among individuals in the host population. The distribution of host susceptibility in the population is quantified by the average transmission rate, 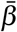, and the *coefficient of variation* in transmission rate among host individuals, *C*. Higher values of *C* mean that there is larger variation in susceptibility among host individuals.

Following well tested theory [25, 43, 45], the effect of including this type of variation in host susceptibility in our model is that individuals with higher susceptibility generally become infected earlier in the epizootic. Remaining hosts are therefore on average more resistant, such that the force of infection 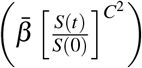 decreases through the epizootic, as the fraction of susceptible hosts remaining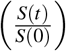 declines. This is the mechanism by which high-resistant hosts can slow down an epizootic and ultimately lead to smaller epizootics (Fig. 4). In addition to determining the number of individuals that become infected and die during an outbreak, the degree of host variation (i.e., the value of *C*) determines whether long-term fluctuations in population size follow point equilibrium dynamics or whether they cycle in a stable or an unstable manner [43].

The epizootic model is linked with a host demographic model. Host reproduction and pathogen over-wintering is described by a set of difference equations, meaning that we assume host reproduction occurs discretely in time [43, 52]. These types of models have been used to describe host-pathogen systems in which the host has seasonal reproduction.

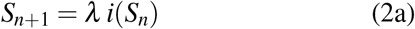

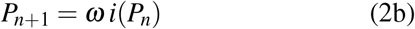

Here, *S*_*n*+1_ and *P*_*n*+1_ are the susceptible host population and pathogen densities in the next generation (*n* + 1). *i*(*S*_*n*_) and *i*(*P*_*n*_) represent the density of susceptible hosts and pathogens that are left after a seasonal epidemic in the current generation, *n*. We assume only the remaining susceptible individuals are available for reproduction. Hosts reproduce at rate *λ*, and the pathogens successfully survive the winter at rate *ω*. Thus the values *S*_*n*+1_ and *P*_*n*+1_ are used as the initial conditions for the next generation’s epidemic (e.g., *S*(0)_*n*+1_ = *S*_*n*+1_ would be the initial condition for equation 1a).

To simulate long-term dynamics of the host-pathogen system, we assume that a seasonal epidemic occurs for 90 days (e.g., during the summer). We therefore simulate the epizootic for 90 days. After 90 days we end the disease epidemic simulation, and we allow the surviving hosts reproduce and pathogens overwinter, following the host demographic model. In the next discrete generation (*n* + 1), overwintered pathogens infect the susceptible available susceptible hosts and initiate a new epidemic. Our model therefore assumes that the effect of the pathogen on long-term host population dynamics is through pathogen-induced mortality during seasonal epidemics. We note that neither the average susceptibility nor the variation in susceptibility are changing across generations due to forces such as selection. The simplified assumption is that environmental conditions exist in each new host generation to generate an equivalent distribution of susceptibilities among host individuals, such that the coefficient of variation in transmission rate, *C*, begins at the same value in each generation.

### Quantifying the effect of temperature

As described above, the amount of variation in host susceptibility (*C*) affects the number of hosts that ultimately become infected during a disease outbreak, and it affects long-term patterns of population cycling. We therefore expect that the level of variation in susceptibility in the host population is a function of temperature (i.e., *C*_(*T*)_, where *T* is temperature), then we would expect the value of *C* to vary seasonally, as average environmental temperatures fluctuate. We hypothesize that allowing *C* to vary through time during an epizootic will result in altered expectations of long-term patterns in host population densities, including altered patterns of cycling.

To test this hypothesis, we used model simulations to characterize population cycles when the variation in host susceptibility shifts during a seasonal epizootic in response to fluctuating environmental temperature. Specifically, we compared two simulations in which we fixed the average susceptibility in the host population 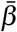, but we allowed two different effects of temperature on the variation in susceptibility, *C*. In the first scenario, we fixed the variation in susceptibility during all seasonal epizootics (Fixed *C, C* = 0.5), meaning there is no effect of temperature. In the second scenario, we allowed variation *C* to depend on environmental temperature (Temperature-Dependent *C*), by assuming that *C* is a function of fluctuating temperature *C*_(*T*)_, and by imposing a temperature regime that generally rises and falls across each 90-day epizootic period (Fig. 5).

**Figure 5:**
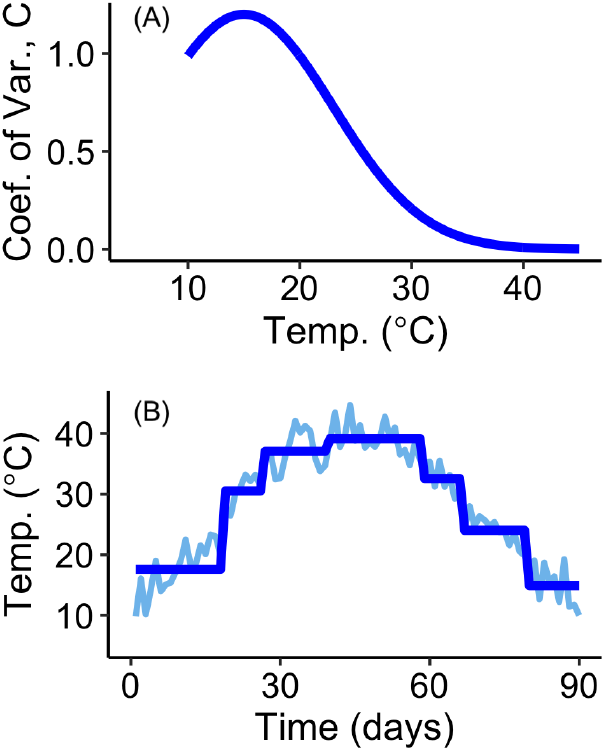
(A) Simulated effect of temperature on *C*, and (B) simulated piece-wise shift in average temperature across the 90-day epizootic.

Because variation *C* is a population-level trait, we assumed that variation *C* gradually responds to the shifting average temperature over time, rather than shifting daily. In other words, we assume that some individuals may respond more quickly than others to temperature, and so observing a measurable difference in *C* may take some time. To reflect this gradual change, we fit a regression tree to simulated temperature data to describe a piece-wise function for changing temperature, from which we calculated a time-specific value of *C*_(*T*)_ (Fig. 5B). In this example, we further assume that variation *C*_(*T*)_ generally declines across a temperature gradient. A pattern of variation (*C*) declining at higher temperatures could arise, for example, if at high temperatures some high-resistance hosts become more susceptible (e.g., due to stress), while some high-susceptibility hosts become more resistant (e.g., improved immune functioning), reducing the overall variance around the mean susceptibility (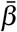). Of course other relationships could be imposed, but we chose a single example to simplify the overall message of our analysis. Finally, to ensure a fair comparison between the two simulated scenarios, we imposed that the average *C*_(*T*)_ over the epizootic was also 0.5 for the Temperature-Dependent *C* scenario.

To simulate the dynamics in the epizootic model, we numerically solved the continuous time differential equations one day forward, replaced the value of *C* to reflect *C*_(*T*)_ at that particular time, and then solved the equations forward another day, continuously for the 90 days. Effectively, this somewhat discretizes the continuous set of differential equations to accommodate the changing values of *C*, following [46].

## Results

When variation is fixed at 0.5 (Fixed *C*), and given our chosen values for the remaining model parameters, the host population exhibits stable cycles, which is what we expect, given previous work [43, 47] (Fig. 6B). However, when the variation in susceptibility among host individuals declines during warmer times of the epizootic (Temperature-Dependent *C*), the cycles in host density are notably different, even though the mean *C* throughout the epizootic was still 0.5 (Fig. 6D). We see that in the Temperature-Dependent *C* scenario there are successive generations of low population numbers, and the population cycles shift from stable to unstable, noticeable around generation 35. This long-term effect on host population stability is attributed to the shifting variation in susceptibility during the epizootic periods. Because temperature reduces variation in susceptibility, the host population temporarily loses some high-resistance individuals (i.e., these individuals become more susceptible). Without these resistant individuals, epizootics can become measurably larger on average, especially at high host densities. When the host population is dense, more hosts get infected and die, causing more severe population crashes after peak host densities. Thus, the host population remains at low densities for successive generations while it recovers from a larger decline after peak host densities, and over time, the population cycles become destabilized.

**Figure 6:**
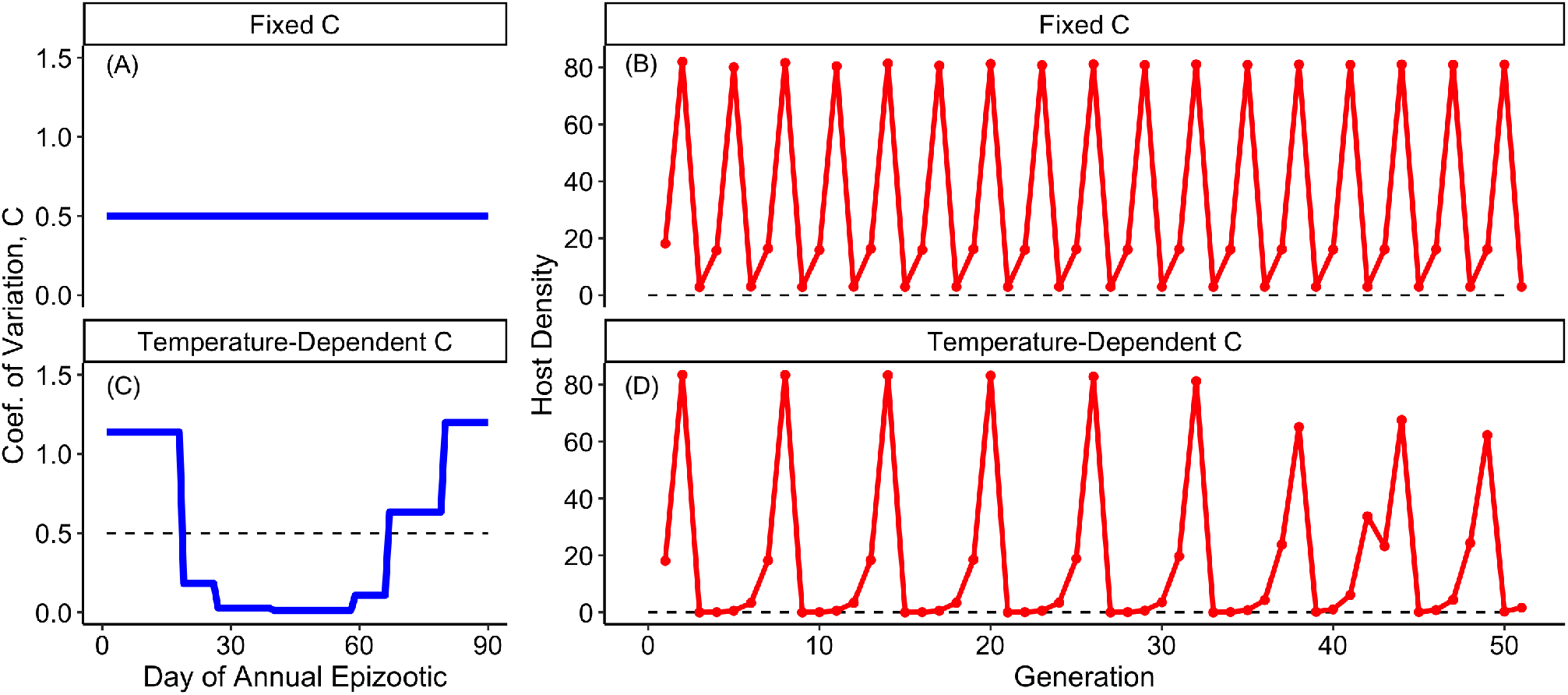
Behavior of host population cycling in response to fixed (A and B) or dynamic (C and D) relationships between variation in susceptibility, *C*, and temperature shifts during epizootics. The dashed line in (C) shows that we fixed the mean *C* = 0.5 to match the mean in (A).

## Discussion

Our model analysis suggests that when among-host trait variation shifts systematically across a thermal gradient, we should expect epizootic patterns to deviate from previous theoretical expectations, and, correspondingly, we should expect unique patterns of cycling host abundances. In general, then, our analysis suggests that the relationship between temperature and among-host variation in traits is an important, but overlooked, mechanism by which climate can mediate short- and long-term disease dynamics. Specifically, this finding has several theoretical and practical implications.

Host-pathogen theory shows that when variation in susceptibility (or more generally, infection risk) is environmentally determined and fixed across generations, we expect a stable point equilibrium of host abundance across generations when *C* > 1; however, when *C* < 1, we expect cycling dynamics (either stable or unstable) [53, 43]. Yet many host populations still cycle when their empirical estimates of *C* are greater than 1, challenging this foundational theory [47]. In such cases, eco-evolutionary dynamics have been proposed as a mechanism by which variation in susceptibility can be realistically high (*C* > 1) and the host population can still cycle. Specifically, if host susceptibility is heritable, under selection, and is costly to fecundity, then stable cycling can still occur when *C* > 1 [47, 49, 45, 42]. Although this eco-evolutionary theory has some empirical support, our analysis introduces an alternative hypothesis: while *C* may be greater than 1 during parts of the epizootic, if temperature drives *C* lower than 1 during other parts, we may still expect cycling. In other words, it may be important to know not just a time-averaged value of *C* over the epizootic, but the specific values *C* may take during an epizootic, if it fluctuates. More generally, a temperature-dependent *C*_(*T*)_ may drive host population cycling in cases that cycling would otherwise be unexpected, although more rigorous theoretical exploration and empirical support of this finding is required.

The short-term effects of temperature on trait variation could also play a major, unappreciated role in determining the threat of pathogens to host population stability in a changing climate. In our analysis, we saw that a temperature-dependent *C*_(*T*)_ could result in larger epizootics, leading to successive host generations of low population abundance, as well as destabilized cycling. As the climate becomes warmer and more variable, the relationship between temperature and trait variation could be a strong predictor of how likely a pathogen is to drive a host population to extinction. Temperature-dependent host variation is therefore an unexplored mechanism of host population declines.

Our results underscore the importance of characterizing the distribution of transmission-related traits among host individuals (e.g., in terms of the average susceptibility *and* the variation in susceptibility), as this will provide fundamental insights into the impact of pathogens on host populations under climate change. Our models are a simplification of the transmission process because temperature is likely to affect the average and the variation in transmission rate, in addition to other parameters that drive epidemic dynamics. We focused on the variation in transmission among host individuals because effects of such variation are mostly omitted from our understanding of disease ecology and of host-pathogen interactions under climate change. Building on the theory presented here is therefore necessary to fully quantify the regulation of disease dynamics by climatic processes.

In general, there has been limited research on the long-term consequences of temperature-dependent traits on host population dynamics in ectotherms. The unpredictability of weather conditions wrought by climate change and the fact that our results show that short-term effects of temperature on host variation can affect long-term population dynamics however suggest that understanding the nature of variation in transmission rates should be a priority of future research. Empirical tests that evaluate the effects of temperatures on transmission rate variability and the variability of other key host and pathogen traits will provide valuable insights into how transmission parameters vary with climate among different taxa. This will prove essential to characterize the potential effects of climate change on host and pathogen populations of concern.

Explicitly measuring effects of temperature on the variation of host traits in a population requires measuring thermal performance curves with adequate replication at each temperature. In addition, studies should ensure that the hosts (or vectors) used in experimentation are a representative sample from the population to capture natural variation. Moreover, it will be important to determine how quickly trait variation among hosts responds to fluctuating temperatures, so that such lags can be adequately incorporated into modeling studies [19]. For some host-pathogen interactions, a close estimate of transmission can be obtained via experimental procedures such as dose-response experiments, where both pathogen dosage and temperature conditions are varied. Models specifically built to estimate transmission parameters can then be applied to these data [25, 54]. However, estimating the effects of temperature on transmission parameters becomes more challenging where experimentation is unfeasible or impractical. In these cases, additional methods exist to statistically infer transmission parameters from time-series data collected from lab-based or naturally-occurring epidemics [55, 56, 46]. These studies could also help reveal whether the effects of temperature-dependent trait averages and variances can be detected at large spatial and temporal scales or whether measurements made at smaller scales (e.g., via experimentation) have meaningful effects on large-scale epidemics and long-term host population dynamics in nature.

To further explore the effect of temperature-dependent variation, modeling studies will need to consider how temperature affects the mean and variance of multiple traits simultaneously. And we need to evaluate in which cases temperature-dependent means or variation will be more important for system dynamics. This poses technical and conceptual challenges. First, to capture inter-individual variation, modeling methods must allow individual hosts (and vectors) to have unique responses to temperature, while imposing any known functional relationships between temperature and the averages and variation of specific traits. This can sometimes be done using carefully designed differential equation models (e.g., Cator et al. [18]), however, it may necessitate building more complex, individual-based models. In constructing complex models, it will be important to evaluate model parsimony, so that general insights can still be derived. Further, sensitivity analysis could be used to compare the effect sizes of temperature-dependent means and variation of certain traits to reveal which traits should be of particular interest for specific systems.

Overlooking the effect of systemic changes in the variation of transmission parameters with climatic variables may provide erroneous predictions about the impact of disease on host population dynamics. Accounting for changes in the distribution of transmission parameters (and not only their averages) among host individuals across environmental gradients and time is however possible through experimentation and the implementation of novel statistical techniques. The application of these methods could make direct contributions to understanding risks of extinction of wildlife populations threatened by climate change.

## Acknowledgements

n/a

## Funding

This material is based upon work supported by the National Science Foundation under Grant No. 2131234. This work was also supported by the State of Arizona Technology and Research Initiative Fund (TRIF), administered by the Arizona Board of Regents, through Northern Arizona University.

## Data, code, and materials

All data and code to run simulations and generate the figures are publicly available on GitHub (https://github.com/joseph-mihaljevic/climate-trait-var)

## Notes

https://github.com/joseph-mihaljevic/climate-trait-var

### Competing Interest Statement

The authors have declared no competing interest.

### Summary of Updates

The manuscript has been heavily reformatted to match a more conventional research article. Figures have been modified to be more piece-meal. Descriptions of the model have been moved from figure to main text.

